# Pregnancy reprograms the enhancer landscape of mammary epithelial cells and alters the response to cMYC-driven oncogenesis

**DOI:** 10.1101/642330

**Authors:** Mary J. Feigman, Matthew A. Moss, Chen Chen, Samantha L. Cyrill, Michael Ciccone, Wesley D. Frey, Shih Ting Yang, John Erby Wilkinson, Camila O. dos Santos

## Abstract

Pregnancy leaves a series of cellular and molecular modifications on mammary epithelial cells (MECs). Pregnancy is also known for decreasing the predisposition of rodent and human MECs to oncogenesis. Here, in order to understand the molecular basis for this effect, we analyzed epigenetic changes in the enhancer landscape of murine post-pregnancy MECs, together with their effect on gene regulation, tissue development and oncogenesis. Using *in vivo* and *in vitro* analyses, we found that completion of a pregnancy cycle changed the dynamics of cellular proliferation and gene expression in response to a second pregnancy. Our results also demonstrated that post-pregnancy MECs are resistant to the initial molecular programs driven by *cMYC* overexpression, a response that blocked MEC proliferation but did not perturb the pregnancy-induced epigenomic landscape. Overall, our findings suggest that pregnancy-induced mammary cancer prevention involves the epigenomic changes in MECs brought about by pregnancy.

## Introduction

In mammals, the process of reproduction is accompanied by substantial developmental reorganization, in response to physiological stimuli from pregnancy. As a result, mammary glands have evolved to assume a pivotal role in milk production and offspring nourishment. Substantial tissue remodeling, including expansion of epithelial cells and ductal structures, is followed by the accumulation of milk droplets as pregnancy progresses. During lactation, milk production is synchronized with milk release by a series of transcriptional and mechanical events present in luminal cells (milk production) and myoepithelial cells (duct contractibility). As lactation ceases, the mammary gland returns to a non-secretory state, and adopts a tissue organization that closely resembles the pre-pregnancy tissue [1].

However, post-pregnancy mammary epithelial cells (MECs) are substantially different from their pre-pregnancy counterparts. Studies have identified transcription networks differentially regulated in post-pregnancy mammary tissue [2–6], showing that it has distinct molecular characteristics compared to the mammary tissue of nulliparous (never pregnant) female mice and humans [7]. Several reports have also suggested that post-pregnancy mammary glands from several mammalian species respond robustly to the signals of consecutive pregnancies [8–11], suggesting a molecular memory of prior pregnancies.

The concept that pregnancy induces long-lasting molecular alterations was also suggested in studies utilizing whole-genome bisulfite sequencing, which showed that pregnancy induced stable and specific losses of DNA methylation in MECs. These epigenetic alterations correlated with the enhanced kinetics of gene re-activation in a subsequent pregnancy [12], suggesting that loss of DNA methylation may underlie epigenetic memory in the post-pregnancy mammary gland.

Pregnancy signals exert functions that go beyond the secretory state, and also modulate the risk of breast cancer in rodents and humans [13–15]. While research has shown increased breast cancer risk for roughly five-ten years after parturition [16–18], there is a long-term reduction of risk of breast cancer for women completing a full-term pregnancy before the age of 30 [13, 14, 18, 19]. A similar risk decrease following pregnancy has been observed in mice, where completion of a pregnancy cycle dampens the frequency of mammary tumor development [14, 20–22]. Given the stability of the molecular programs brought by pregnancy to MECs, and the longevity of cancer preventive effects in rodents and in humans, it is possible that such effects have an epigenetic basis.

To address this hypothesis, we set out to characterize the dynamics of gene reactivation and enhancer activity in murine MECs as they respond to pregnancy signals or early oncogenesis. Global gene expression analysis of active regulatory regions (H3K27ac ChIP-seq) revealed epigenomic alterations established early in the first pregnancy cycle, which influenced the transcription output of post-pregnancy MECs when re-exposed to pregnancy signals (second pregnancy). This rapid response to consecutive pregnancy cycles was preserved even after the transplantation of post-pregnancy MECs into the cleared fat pad of virgin female mice, supporting our hypothesis that pregnancy stably alters the epigenome of MECs.

To characterize the effects of a pregnancy-induced epigenome in response to oncogenic stress, we established a transgenic mouse strain (CAGMYC), in which overexpression of the oncogene *cMYC*, a known inducer of mammary tumor development [7, 23, 24], is driven in a doxycycline-dependent manner. Using this transgenic mouse strain, we found that the post-pregnancy epigenome was incompatible with *cMYC* overexpression, blocking the activation of MYC-downstream signals and their progression to oncogenesis.

Overall, our findings suggest that pregnancy-induced mammary cancer prevention involves the epigenomic changes in MECs brought about by pregnancy, which now can be leveraged to guide studies investigating the effects of pregnancy on breast cancer risk in human tissue.

## Results

### Pregnancy-induced epigenome in post-pregnancy MECs

The observation that pregnancy induces loss of DNA methylation levels at specific genomic regions in post-pregnancy MECs suggests that such regions assume an active regulatory state after pregnancy [12]. To test this idea, we mapped global gene expression (RNA-seq) of FACS-isolated MECs from nulliparous (pre-pregnancy) and parous (post-pregnancy) female mice, as well those harvested from female mice during the first and second pregnancy. For the first and second pregnancy time points, nulliparous Balb/c female mice, or Balb/c female mice that completed one natural pregnancy, were treated with slow-release estrogen and progesterone hormones to simulate pregnancy, to ensure a precise timing of pregnancy hormone exposure. This procedure induces mammary gland development that closely resembled both the histological and epigenetic modifications observed in mice where pregnancy occurred following conception [25] (Supplementary Fig. 1a).

Pre- and post-pregnancy MECs had similar transcription programs, as revealed by unsupervised expression clustering, suggesting that their epithelial identity during tissue homeostasis is not substantially altered by a previous pregnancy cycle. MECs harvested during the early stages of a second pregnancy (D6) clustered together with those harvested at a later time-point during a first pregnancy (D12), supporting that post-pregnancy MECs respond robustly to consecutive pregnancy signals (Fig.1a). In order to address whether this robust response to second pregnancy signals have an epigenetic basis, we investigated the genomic distribution of the active histone mark H3K27ac, which annotates active regulatory elements (promoters and enhancers) in mammalian genomes, in the same cohort of MECs utilized for the RNA-seq analysis (Supplementary Fig. 1a).

**Figure 1.**
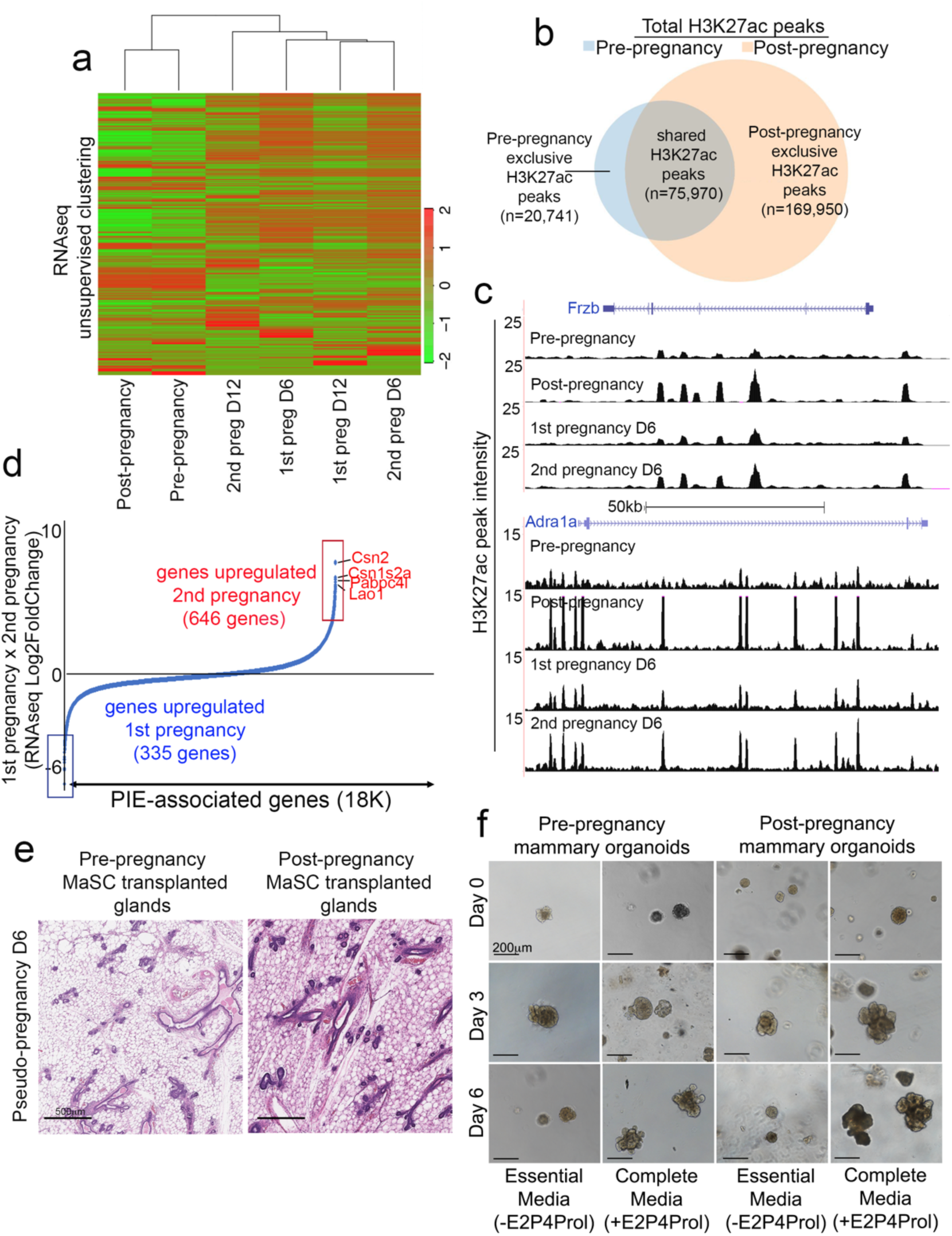
Pregnancy-induced epigenome in post-pregnancy MECs. (**a**) Heatmap distribution of gene expression data collected from FACS-isolated MECs spanning several developmental stages. (**b**) Venn-diagram demonstrating the number of shared and exclusive H3K27ac ChIP-seq peaks of FACS-isolated MECs from pre-pregnancy female mice (blue circle) and post-pregnancy female mice (orange circle). (**c-**) Genome browser tracks showing distribution of H3K27ac peaks at distinct pregnancy cycles for Frzb and Adra1a loci. (**d**) Expression of genes associated with Parity Induced Elements (PIEs), according to Log2 fold change (differential expression) in luminal MECs harvested from mammary glands during first and second pregnancy. Boxes indicate genes most abundantly upregulated during first (log2FoldChange ←­2) (blue) and second pregnancy (log2FoldChange >= 2) (red). (**e**) H&E stained mammary glands images, from glands transplanted with either pre-pregnancy (left panel) or post-pregnancy (right panel) MaSCs, harvested on day 6 of pseudo-pregnancy. Images were acquired on Aperio ePathology (Leica Biosystems) slide scanner. Scale: 500 µm. (**f**) 3D Matrigel™ mammary organoid culture of pre- and post-pregnancy MECs, grown with either Essential media or Complete media (containing Estrogen (E2), Progesterone (P4) and prolactin (Prol)). Images captured on Nikon Eclipse TI microscope and processed on NIS-Elements BR (Nikon). Scale: 200 µm.

Post-pregnancy MECS exhibited an eight-fold increase in the number of exclusive H3K27ac peaks (n=169,950), compared to pre-pregnancy MECs (n=20,741), demonstrating that pregnancy significantly expands the regulatory landscape of MECs (Fig. 1b). H3K27ac peaks in pre-pregnancy MECs were enriched in pathways associated with kinase activity, RNA binding and stem cells (Supplementary Fig. 1b), while in post-pregnancy MECs, the H3K27ac peaks enriched for pathways involved in regulation of cell polarity, mRNA splicing, DNA methylation and response to stress (Supplementary Fig. 1c). These observations show that pre- and post-pregnancy MECs engage in distinct cellular maintenance and tissue homeostasis pathways.

Further investigation of the regulatory regions demarcated by the pregnancy-induced epigenome demonstrated that the majority of H3K27ac peaks appearing after pregnancy resided within genic and intergenic regions, likely representing distal enhancer elements (Supplementary Fig. 1d), suggesting that pregnancy-induced epigenomic changes may specifically expand the enhancer landscape of MECs. To identify H3K27ac peaks at enhancer/super enhancer regions and probe their role in gene regulation, we utilized ROSE [26] to stitch together nearby peaks located at genic and intergenic regions and thereby delineate enhancer regions. Approximately 5,000 enhancers were defined by H3K27ac peaks in pre-pregnancy MECs (Supplementary Fig. 1e, top panel), in clear contrast to post-pregnancy MECs, which had over 60,000 enhancer regions, referred to hereafter as Parity Induced Elements (PIEs). These results support the notion that pregnancy expands the enhancer landscape of MECs (Supplementary Fig. 1e, bottom panel). Interestingly, a large fraction of PIEs were at an active state (enriched for H3K27ac) during the first pregnancy and during the second pregnancy, further supporting the stability of the pregnancy-altered epigenome (Fig. 1c and Supplementary Fig. 1f).

We utilized this dataset to define genes associated with enhancers that were exclusively present in post-pregnancy MECs, revealing ∼18,000 PIE-associated genes. These PIE-associated genes were further utilized to define the dynamics of the pregnancy-induced epigenome, and their effects on the transcriptome, across pregnancy cycles (first pregnancy and second pregnancy). Our analysis revealed that over 600 genes associated with PIEs were more abundantly expressed in MECs harvested from mice during the second pregnancy cycle, with the top hits being casein-associated genes, known to have roles in milk production [27], which is supportive of a robust activation of pregnancy-related programs (Fig. 1d, red) [12].

It is possible that mammary stem cells (MaSCs) and progenitor cells propagate the epigenetic memory of pregnancy. It is also possible that alterations to the post-pregnancy non-epithelial cells perpetuate this epigenetic state, thus locking post-pregnancy MECs into a pregnancy-prone state. In order to dissect the role of the microenvironment on the ability of post-pregnancy MECs to respond to a second pregnancy, we employed fat-pad transplantation assays [28, 29]. Mammary glands from pre-pubescent, virgin female mice (15-days old) were cleared of endogenous epithelium and transplanted with either pre- or post-pregnancy MaSCs. Recipient female mice (2-months post-transplantation) were then exposed to pseudo-pregnancy, followed by histology analysis of mammary glands (Supplementary Figure 1 g).

Transplanted mammary glands from placebo-treated females demonstrated comparable ratios of luminal and myoepithelial cells to glands that received either pre- or post-pregnancy MaSCs, suggesting that pregnancy did not change the ability of transplanted MaSCs to undergo lineage commitment and differentiation (Supplementary Fig. 1 h). Moreover, RT-qPCR analysis of defined genes up-regulated after pregnancy demonstrated that despite transplantation into a pregnancy-naïve environment, post-pregnancy MECs retained the ability to regulate pregnancy-induced genes (Supplementary Fig. 1i).

Tissue histology analysis during pseudo-pregnancy demonstrated that mammary glands transplanted with post-pregnancy MaSCs had higher numbers of ductal structures (<u>668 ducts,+/- 32)</u> than glands transplanted with pre-pregnancy MaSCs (467 ducts, +/- 93), suggesting that fat-pad transplantation does not alter the ability of post-pregnancy MECs to react robustly to pregnancy signals (Fig. 1e; Supplementary Fig. 1j, p<0.05). This rapid response to pregnancy hormones was also recapitulated using an organoid culture system, where we found that organoids derived from post-pregnancy MECs displayed increased branching morphogenesis (microscope analysis) and gene expression levels (qPCR analysis), than those cultures derived from pre-pregnancy MECs (Fig. 1f and Supplementary Figure 1k). Thus, our findings support that intrinsic, epigenetic-based, cell-autonomous signals direct post-pregnancy MECs to respond more rapidly and efficiently when re-exposed to the signals of pregnancy.

### *cMYC* drives epigenomic and transcriptomic alterations during mammary oncogenesis in pre-pregnancy MECs

Pregnancy decreases the frequency of mammary tumors in a variety of transgenic and carcinogenic mouse models of mammary oncogenesis [13–15], raising the possibility that pregnancy-induced epigenomic and transcriptomic modifications may underlie pregnancy-induced mammary cancer protection. This led us to hypothesize that the enhancer reprogramming observed after pregnancy might influence the output of oncoprotein that function via a transcriptional mechanism. Several of the above-mentioned studies have utilized transgenic mice in which oncogenes were under control of mammary gland-specific promoters, such as MMTV and WAP-CRE, to drive tumor development. However, such promoters have been shown to be enhanced by several signals present during pregnancy and lactation [30–32], which could mask a clear understanding of the epigenomic and transcriptomic changes associated with early oncogenesis and pregnancy-induced protection. To overcome this potential obstacle, we used a mouse model in which overexpression of the potent oncogene cMYC was controlled by a tetracycline regulatory system (Doxycycline induced, DOX), under the control of the CAG promoter (independent of pregnancy/lactation signals), referred to hereafter as CAGMYC (Supplementary Fig. 2a).

We found that sustained DOX administration through drinking water for 2 days (DD2) or 5 days (DD5) resulted in substantial morphological alterations to the mammary gland of CAGMYC transgenic nulliparous female mice (Fig. 2a), including tissue hyperplasia, with expansion and flattening of ductal structures, frequently observed during the initial stages of murine mammary oncogenesis [33] (Fig. 2a). None of these alterations were seen in control CAG-only transgenic mice (Supplementary Fig. 2b), thus indicating that short-term *cMYC* overexpression is sufficient to drive hyperplasia of mammary glands from nulliparous mice.

**Figure 2.**
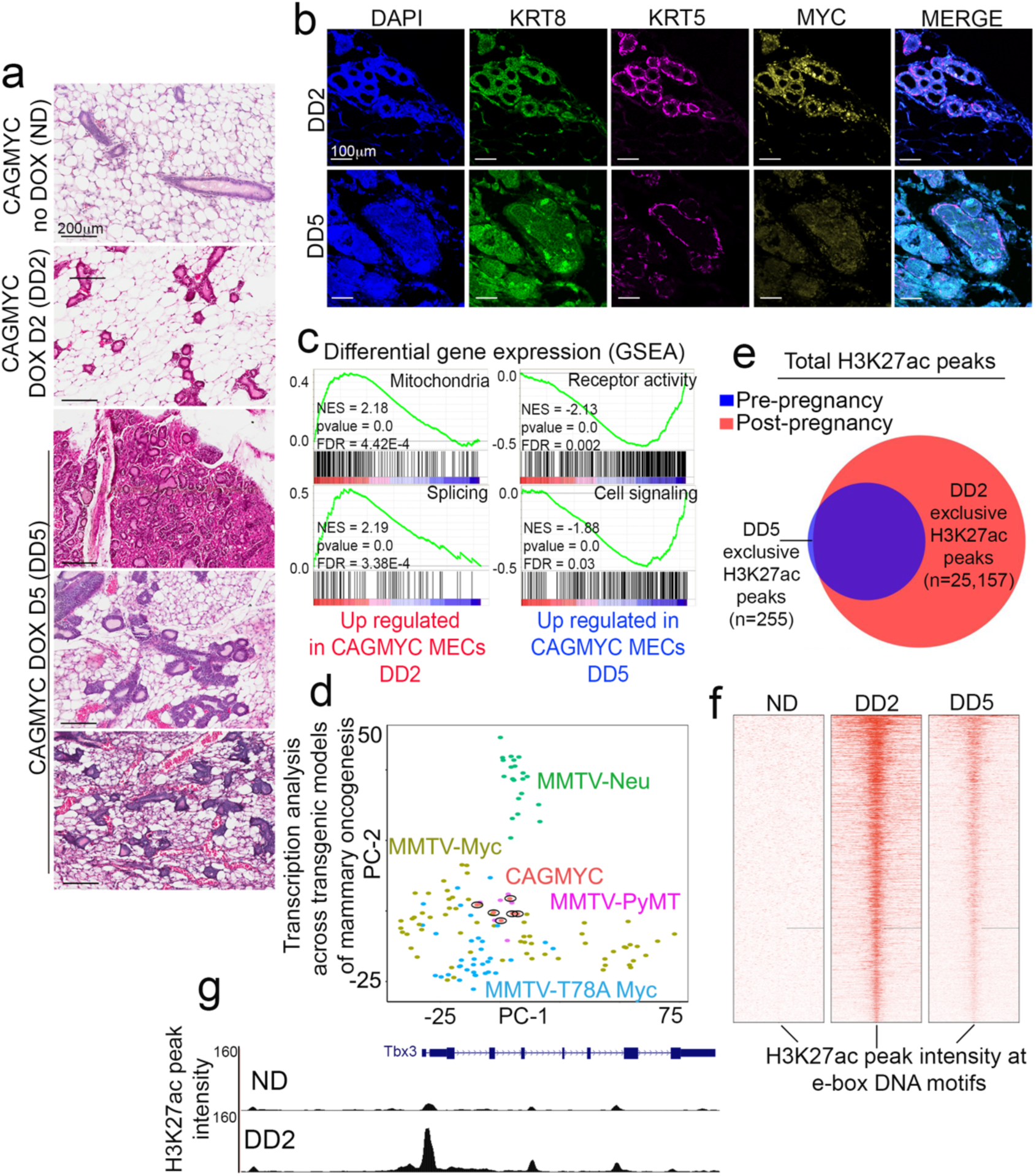
*cMYC* drives epigenomic and transcriptomic alterations during mammary oncogenesis in pre-pregnancy MECs. (**a**) H&E mammary glands images from virgin CAGMYC female mice, untreated (ND) and treated with DOX for 2 days (DD2) and for 5 days (DD5). Scale: 200 µm. (**b**) Immunofluorescence images of mammary glands from nulliparous CAGMYC female mice treated with DOX for 2 (DD2) and 5 days (DD5), visualizing DAPI (blue), KRT8 (green), KRT5 (magenta) and cMYC (yellow). Images captured on Zeiss LSM780 laser scanning microscope and processed on Zen lite (Zeiss). Scale: 100 µm. (**c**) GSEA analysis for enriched transcriptional programs CAGMYC MECs harvested from virgin female mice treated with DOX for 2 days (DD2) and for 5 days (DD5). NES: Normalized Enrichment Score. (**d**) Principal Component analysis (PCA) of gene expression levels from CAGMYC MECs harvested from virgin female mice treated with DOX, compared to publicly available gene expression datasets from additional transgenic mouse models of mammary tumorigenesis. (**e**) Venn-diagram demonstrating the number of shared and exclusive H3K27ac ChIP-seq peaks of CAGMYC MECs harvested from virgin female mice treated with DOX for 2 days (DD2, red) and for 5 days (DD5, blue). (**f**) Heatmap of distribution of H3K27ac peaks at e-box DNA binding motifs, in MECs harvested from virgin female mice untreated (ND), treated with DOX for 2 (DD2) and 5 days (DD5). (**g**) Genome Browser tracks for Tbx3 gene, showing differential H3K27ac distributions in CAGMYC MECs upon cMYC overexpression.

To study the alterations to mammary glands in DOX-treated CAGMYC female mice, we analyzed their cytokeratin composition. We observed a progressive expansion of cytokeratin 8 (KRT8) expressing cells, a hallmark for luminal-like cells [34], over the course of the DOX treatment (Fig. 2b). This phenotype was accompanied by progressive thinning of the cellular layer that constitutes basal-like cells (cytokeratin 5, KRT5), often observed during mammary tissue hyperplasia, and that precedes the extravasation of luminal cells from ductal structures [35] (Fig. 2b). In fact, DD5 mammary glands showed extravasation of KRT8+ cells outside KRT5+ cellular boundaries, further supporting that *cMYC* overexpression drives hyperplasia in the mammary tissue.

Given that the cMYC-driven mammary tumors have been shown to drive pathologically and transcriptionally heterogeneous mammary tumors [36], we asked whether the transcriptome of CAGMYC MECs would shed light on their mammary cancer subtype classification. CAGMYC MECs were FACS-isolated from transgenic mice during sustained *cMYC* overexpression (DD2 and DD5) and analyzed using RNA-seq. In DD2 mice we observed up-regulation of pathways involving cellular metabolism, such as mitochondrial function and gene splicing. In contrast, pathways up-regulated in DD5 mice were associated with control of cell communication processes (Fig. 2c). These results suggest that transcriptional programs controlled by *cMYC* overexpression progressively alter MECs, as part of the global oncogenesis program. These observations also corroborate our histological analysis revealing progressive alterations to mammary gland architecture in response to sustained *cMYC* overexpression (Fig. 2a, b). Collectively, these findings validate the CAGMYC transgenic mouse system as a suitable model for investigating ductal hyperplasia and the initial stages of murine mammary oncogenesis.

To gain a deeper understanding of the lineage profile of CAGMYC MECs, we compared DD2 and DD5 CAGMYC MEC gene expression patterns across publicly available transcription datasets for various transgenic models of mammary oncogenesis [37]. We found that the transcriptional profiles of CAGMYC MECs clustered closely with those from mixed and luminal-like mammary tumors, including those harvested from MMTV-Myc models (Fig. 2d, Supplementary Fig. 2c). Tissue staining with antibodies against estrogen receptor alpha (ERα), a hallmark of the most common luminal-like tumor subtypes, demonstrated that only a subset of DOX-treated CAGMYC MECs expressed the receptor (Supplementary Fig. 2d). Therefore, these results support that the ductal hyperplasia observed in CAGMYC transgenic mice carry a heterogeneous, mixed, luminal-biased cellular phenotype that resemble those observed in other *cMYC* transgenic lines of mammary oncogenesis [36][37].

We next investigated the effects of short-term *cMYC* overexpression on the enhancer landscape of MECs by mapping the active H3K27ac enhancer landscape (H3K27ac ChIP-seq) of CAGMYC MECs. Our analysis demonstrated only a slight gain of H3K27ac peaks in DD5 versus DD2 CAGMYC MECs, suggesting that many regions activated by *cMYC* overexpression at DD5 were already altered at DD2 (Fig. 2e). Genomic distribution analysis showed that this gain of H3K27ac occurred mainly at promoter regions (Supplementary Fig. 2e), an observation that agrees with previous reports demonstrating that *cMYC* induced chromatin activation occurs mainly at promoter regions [38]. Gene ontology analysis revealed that many H3K27ac peaks induced after *cMYC* overexpression at DD2 or DD5 mapped to genes controlling signals associated with increased proliferation, transcription and signaling (Supplementary Fig. 2f), further suggesting that *cMYC* overexpression induces signals that support abnormal mammary hyperplasia.

A prior study demonstrated that *cMYC* deletion in mice impaired ductal alveolar genesis during pubescence and pregnancy, indicating it is required for normal mammary gland development [39]. Thus, we asked whether *cMYC* overexpression activated a distinct enhancer landscape compared to that seen during normal development. We focused on the gain of H3K27ac signal at genomic regions preferentially recognized by cMYC (e-boxes) [40] in response to *cMYC* overexpression. We found that approximately 2,500 H3K27ac peaks were detected at e-boxes in wild-type, non-transgenic MECs, suggesting that a defined set of cis-regulatory elements are activated by *cMYC* in normal MECs (Supplementary Fig. 2 g).

In contrast, we detected a gain of H3K27ac peaks at specific e-boxes from CAGMYC MECs in response to *cMYC* overexpression, suggesting activation of transcription networks selective for oncogenesis (Fig. 2f). Genome browser images also show gain of H3K27ac in CAGMYC MECs after induction of *cMYC* overexpression, at known cMYC downstream targets, which have been implicated in mammary oncogenesis [41, 42] (Fig. 2 g, Supplementary Fig. 2 h). Thus, our results demonstrate that short-term *cMYC* overexpression activates a specific transcriptional network, that differs from those regulated during normal development, and drives tissue morphology alterations resembling those of murine mammary ductal hyperplasia.

### *cMYC* fails to drive oncogenesis in post-pregnancy MECs

To investigate the effects of *cMYC* on post-pregnancy CAGMYC MECs, we induced *cMYC* overexpression in parous CAGMYC female mice (post-pregnancy, 21-days gestation, 20-days lactation, 40-days involution) for 5 days (DD5) (Supplementary Fig. 2a). Remarkably, histological analysis revealed that mammary glands of DD5 parous CAGMYC female mice remained largely unaffected by *cMYC* overexpression (276 ducts, +/-100), displaying tissue morphology similar to mammary glands from DOX-treated, CAG-only control groups (387 ducts, +/-10) Fig. 3a, Supplementary Fig. 3b). This was in marked contrast to the control cohort of nulliparous CAGMYC female mice, which instead displayed mammary gland ductal hyperplasia (789 ducts, +/-185), in response to *cMYC* overexpression (Fig. 2a, Fig. 3a, b, Supplementary Fig. 3c, pvalue= 0.0001). Such phenotypic discrepancies were not caused by inefficient transgene induction, as pre- and post-pregnancy CAGMYC MECs expressed comparable *cMYC* mRNA and protein (Fig. 3c, Supplementary Fig. 3d). Thus, post-pregnancy MECs do not respond to early oncogenesis signals driven by *cMYC* overexpression.

**Figure 3.**
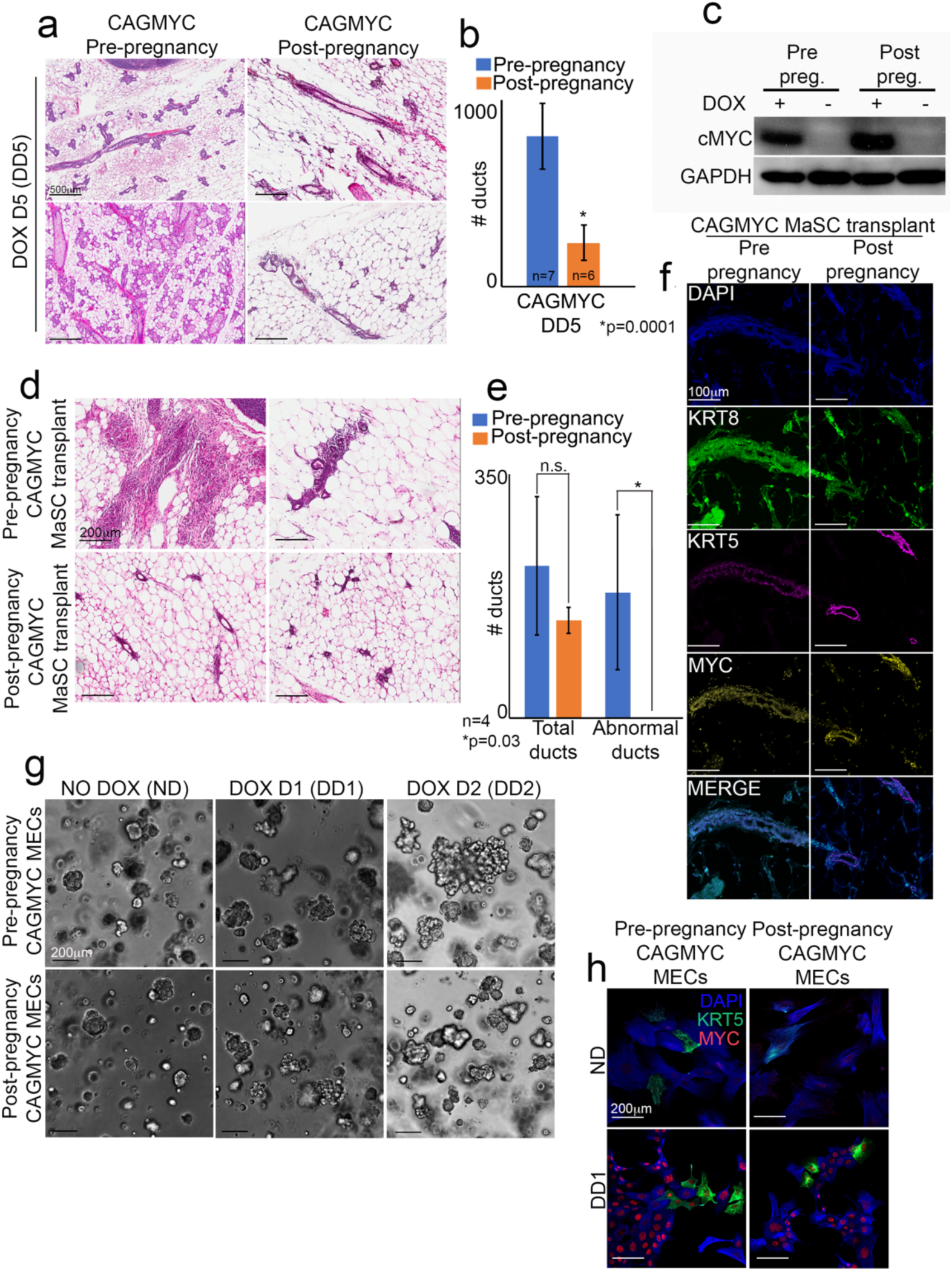
*cMYC* fails to drive oncogenesis in post-pregnancy MECs. (**a**) H&E stained mammary glands images from nulliparous (right panel) and parous (left panel) CAGMYC female mice treated with DOX for 5 days. Images acquired on Aperio ePathology (Leica Biosystems) slide scanner. Scale: 500 µm. (**b**) Quantification of ductal structures in H&E-stained mammary glands from nulliparous (left bar, n=7) and parous (right bar, n=6) CAGMYC female mice treated with DOX for 5 days. Bars indicate mean number of ducts. Error bars indicate standard deviation amongst samples of same experimental group. *pvalue= 0.0001. (**c**) Western blot of cMYC protein (62 kDa) in pre-pregnancy and post-pregnancy CAGMYC MECs, with and without DOX treatment (5 days). GAPDH (146 kDa) used as endogenous control. (**d**) H&E stained mammary glands images from nulliparous CAG-only control mice, transplanted with pre-pregnancy and post-pregnancy CAGMYC MaSCs and treated with DOX for 5 days. Scale: 200 µm. (**e**) Quantification of ductal structures from H&E stained images of mammary glands from nulliparous CAG-only control mice, transplanted with pre-pregnancy (n=4) and post-pregnancy (n=4) CAGMYC MaSCs and treated with DOX for 5 days. Bars indicate mean number of total ducts (left) and abnormal ducts (right). Error bars indicate standard deviation amongst samples of same experimental group. n.s. = not significant. *pvalue = 0.003. (**f**) Immunofluorescence images of mammary glands from nulliparous CAG-only control mice, transplanted with pre-pregnancy and post-pregnancy CAGMYC MaSCs and treated with DOX for 5 days, visualizing DAPI (blue), KRT8 (green), KRT5 (magenta) and cMYC (yellow). Images captured on Zeiss LSM780 laser scanning microscope and processed on Zen lite (Zeiss). Scale: 100 µm. (**g**) 3D Matrigel™ mammary organoid culture of pre- and post-pregnancy CAGMYC MECs, grown with either Essential media and with or without DOX (0.1 mg/mL). Images captured on Nikon Eclipse TI microscope and processed on NIS-Elements BR (Nikon). Scale: 200 µm. (**h**) Immunofluorescence image of pre-pregnancy and post-pregnancy CAGMYC MECs harvested from organoid cultures and treated with DOX for 24 hours, visualizing DAPI (blue), KRT5 (green), cMYC (yellow). Images captured on Zeiss LSM780 laser scanning microscope and processed on Zen lite (Zeiss). Magnification: 20X. Scale: 200 µm.

In prior studies, pregnancy-induced mammary cancer protection was studied in rodents in which pregnancy signals were mimicked by the implantation of hormone pellets (pseudo-pregnancy) [43, 44]. Thus, we tested whether pseudo-pregnancy would block the oncogenic effect of *cMYC* overexpression on MECs. Nulliparous CAGMYC female mice were implanted with slow-release estrogen and progesterone pellets and analyzed 60-days post-pellet implantation (∼20 days pseudo-pregnancy and ∼40 days of forced involution), followed by DOX treatment (DD5) (Supplementary Fig. 3e). All post pseudo-pregnancy mammary glands analyzed (n=3) displayed abnormal tissue morphology, marked by tissue hyperplasia resembling the finals stages of involution [45, 46] (Supplementary Fig. 3f, bottom panel), unlike the phenotype observed in nulliparous CAGMYC female mammary glands (Fig. 2a, Fig. 3a). Analysis of pseudo-pregnancy in CAG-only control transgenic mice revealed a similar mammary gland phenotype as observed in CAGMYC transgenic mice (Supplementary Fig. 3f, top panel), suggesting that abnormal tissue histology in pseudo-pregnancy-induced mice was not a result of *cMYC* overexpression. Given that mammary gland tissue abnormalities have been reported to occur in a strain-specific manner in response to slow-release pellet implants [47], we next performed histologic analysis in C57/BL6 and Balb/C female mice during pseudo-pregnancy. We found that completion of pseudo-pregnancy in C57/BL6 and Balb/C mouse strains revealed substantially fewer ductal structures, similar to the tissue architecture of a normal, fully involuted mammary gland (Supplementary Fig. 3 g), and in marked contrast to the phenotype observed in CAG-only transgenic mice (Supplementary Fig. 3f). Thus, the post pseudo-pregnancy abnormalities in mammary glands from transgenic CAGMYC and CAG-only mice are likely a consequence of its genetic strain. Given this abnormal morphology, we focused our analysis of ductal hyperplasia in CAGMYC MECs using female mice exposed to natural pregnancy cycles (gestation, lactation and full involution).

To explore whether the ability of post-pregnancy CAGMYC MECs to resist malignancy is driven by cell-autonomous or non-autonomous mechanisms, pre- and post-pregnancy CAGMYC MaSCs were transplanted into the fat-pad of CAG-only, nulliparous female mice, followed by DOX treatment for 5 days (Supplementary Fig. 3 h). We used transgenic CAG-only female mice as hosts for the transplantation assays to avoid immune rejection driven by CAGMYC MaSCs transgene expression [48]. Transplantation of post-pregnancy CAGMYC MaSCs did not result in the development of abnormal ductal clusters in response to *cMYC* overexpression (p=0.03), in contrast to fat-pads transplanted with pre-pregnancy CAGMYC MaSCs, which displayed hyperplastic ductal clusters (180 abnormal ducts, +/-111, Fig. 3d, e). Glands transplanted with post-pregnancy CAGMYC MaSCs displayed similar numbers of overall ducts (140 ducts, +/-18) compared to those from glands transplanted with pre-pregnancy CAGMYC MaSCs (218 ducts, +/-99), suggesting that a lack of abnormal ductal clusters was not an artifact associated with transplantation (Fig. 3e, f). Given the lack of abnormal morphology of the mammary glands transplanted post-pregnancy CAGMYC MaSCs, our results suggest that pregnancy-induced, cell-autonomous signals block ductal hyperplasia, even when post-pregnancy MaSCs/MECs are exposed to a pregnancy-naïve environment.

To investigate whether the cell-autonomous, hyperplasia-resistance phenotype of post-pregnancy MECs would persist under *in vitro* growth conditions, we utilized a mammary organoid system to test the growth and oncogenic potential of pre- and post-pregnancy CAGMYC MECs. Analysis of pre- and post-pregnancy organoid cultures exposed to increasing concentrations of DOX (0.01 mg/mL, 0.1 mg/mL and 0.5 mg/mL) demonstrated similar induction of *cMYC* overexpression levels, suggesting that DOX-inducible activation of transgene expression is comparable in both conditions (Supplementary Fig. 3e, i). Morphological analysis of DOX-untreated organoid cultures demonstrated that pre- and post-pregnancy CAGMYC organoid cultures displayed similar growth kinetics and branch-like morphology (Fig. 3 g, left panel). DOX treatment (0.1 mg/mL) of organoid cultures, however, resulted in progressive morphological change and growth of the pre-pregnancy CAGMYC organoids (Fig. 3 g), marked by increased cell density in the center of the organoids and increased organoid size, phenotypes not observed in cultures of post-pregnancy CAGMYC organoids (Fig. 3 g). The absence of a phenotypic response to *cMYC* overexpression in post-pregnancy organoids was also observed in cultures supplemented with high concentrations of DOX (Supplementary Fig. 3f, Fig. 3 h). Thus, our results support that even in *in vitro* cultures, post-pregnancy CAGMYC MECs do not respond to the oncogenic effects of *cMYC* overexpression.

### cMYC-driven effects are blocked in CAGMYC post-pregnancy MECs

We next set out to define the transcriptome and epigenome of post-pregnancy CAGMYC MECs, in order to compare them to the *cMYC*-driven oncogenic effects seen in pre-pregnancy CAGMYC MECs (Supplementary Fig.4a). Transcriptome analysis of luminal and myoepithelial cells from pre- and post-pregnancy CAGMYC female mice (DD5) demonstrated that overall *cMYC* overexpression did not alter lineage-specific transcription, given that luminal cell and myoepithelial cells clustered together, independent of parity (Fig. 4a). Despite the unaltered lineage specification, differential gene-expression analysis demonstrated that post-pregnancy CAGMYC MECs failed to activate transcription networks controlled by cMYC, such as canonical cMYC targets, response to estrogen and mTORC1 signaling, thus in agreement with the lack of hyperplasia in mammary tissue from CAGMYC parous mice (Fig. 4b). Networks down-regulated in post-pregnancy CAGMYC MECs included genes essential for mammary development and morphogenesis (Supplementary Fig. 4b), indicating that specific mammary epithelial programs do not respond to cMYC-driven oncogenesis.

**Figure 4.**
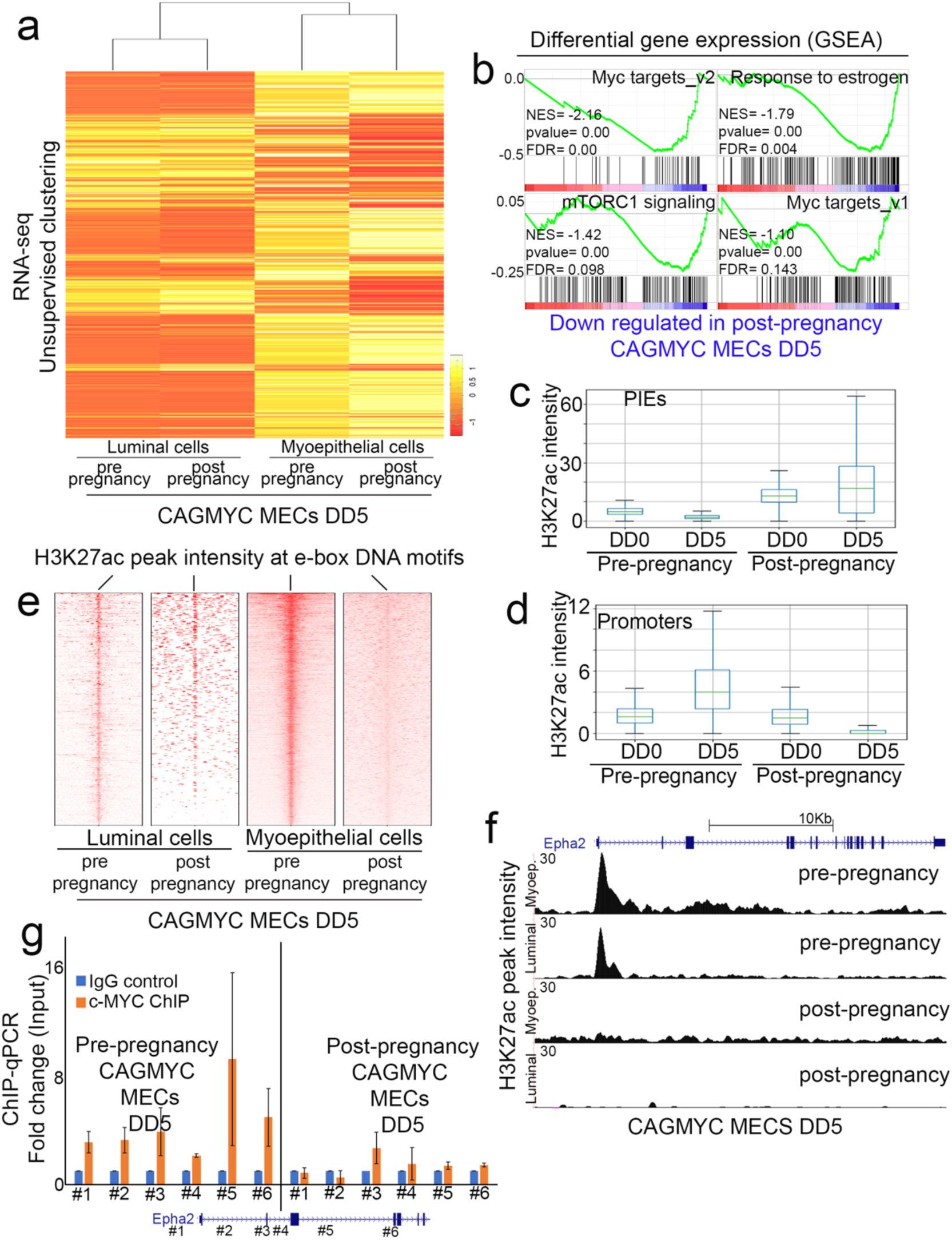
cMYC-driven effects are blocked in CAGMYC post-pregnancy MECs. (**a**) Hierarchical clustering of gene expression of DOX-treated, FACS-isolated, pre- and post-pregnancy CAGMYC MECs (luminal and myoepithelial). (**b**) GSEA analysis of differential RNA expression in DOX-treated, FACS-isolated, pre-(blue) and post-pregnancy (red) CAGMYC MECs. NES: Normalized Enrichment Score. (**c,d**) Analysis of H3K27ac intensity at (**c**) Parity-Induced Elements (PIEs) or (**d**) promoter regions in DOX-treated, FACS-isolated, pre- and post- pregnancy CAGMYC MECs. Box plots display mean averaged H3K27ac intensity and error bars represent the variation of H3K27ac intensity at analyzed regions. (**e**) Heatmaps demonstrating H3K27ac intensity at e-box DNA binding motifs in DOX-treated, FACS-isolated (luminal and myoepithelial), pre- and post-pregnancy CAGMYC MECs. (**f**) Genome Browser tracks for EphA2 gene showing differential H3K27ac distributions in DOX-treated, FACS-isolated (luminal and myoepithelial), pre- and post-pregnancy CAGMYC MECs. (**g**) ChIP-qPCR quantification of cMYC occupancy at the EphA2 gene locus DOX-treated, pre- and post-pregnancy CAGMYC MECs. Bars indicate fold enrichment in relation to Input.

These results suggest post-pregnancy MECs are less responsive to the transcription activation elicited by *cMYC* overexpression. We therefore performed H3K27ac ChIP-seq on pre- and post-pregnancy CAGMYC MECs (DD5) to address the extent of the epigenomic changes in post-pregnancy cells (Supplementary Fig. 4a). Genome distribution analysis revealed that *cMYC* overexpression expanded the number of H3K27ac peaks at promoter regions specifically in pre-pregnancy CAGMYC MECs. Conversely, post-pregnancy CAGMYC MECs were marked by increased H3K27ac peaks at intergenic and genic regions, supporting the notion that pregnancy induces an expansion of the enhancer landscape in MECs, and suggesting that the transcriptional output of cMYC is altered by transitions through pregnancy (as seen in Fig. 1), which is not erased by *cMYC* overexpression (Supplementary Fig. 4c). Further analysis demonstrated that H3K27ac peaks from DOX-induced post-pregnancy CAGMYC MECs overlapped minimally with those detected in pre-pregnancy CAGMYC MECs (Supplementary Fig. 4d). Gene ontology analysis of H3K27ac peaks seen exclusively in post-pregnancy CAGMYC MECs revealed enrichment for pathways that negatively control mammary gland development, suggesting that pregnancy-induced epigenetic mechanisms may also block cellular proliferation in response to *cMYC* overexpression (Supplementary Fig. 4e).

Analysis of PIEs (defined on Fig. 1) demonstrated low H3K27ac intensity in pre-pregnancy CAGMYC MECs, independent of *cMYC* overexpression. Conversely, post-pregnancy CAGMYC MECs displayed a further increase in H3K27ac signal, suggesting that the regions associated with robust responses to pregnancy remained unaltered in response to *cMYC* overexpression (Fig. 4c). In agreement, *cMYC* overexpression specifically increased the intensity of H3K27ac peaks at promoter regions in pre-pregnancy CAGMYC MECs, further suggesting that signals driven by cMYC preferentially activate promoter regions in MECs, which are not activated in post-pregnancy CAGMYC MECs (Fig. 4d).

Given that cMYC preferentially occupies e-boxes (Fig. 2), we asked whether these would be differently enriched for H3K27ac signals in pre- and post-pregnancy CAGMYC MECs. This analysis should also remove any promoter or enhancer bias in the H3K27ac signal in pre- and post-pregnancy CAGMYC MECs (Fig. 4c, d). These analyses revealed that both post-pregnancy CAGMYC luminal and myoepithelial MECs failed to accumulate H3K27ac peaks at e-boxes (∼5,000 regions) in response to *cMYC* overexpression (Fig. 4e). This was in marked contrast to the increased H3K27ac signal intensity detected at the same regions in pre-pregnancy CAGMYC MECs, further supporting the notion that post-pregnancy CAGMYC MECs were insensitive to gene activation from *cMYC* overexpression (Fig. 4e).

Several of the genes down-regulated in post-pregnancy CAGMYC MECs displayed substantially lower levels of H3K27ac and cMYC DNA occupancy after *cMYC* overexpression (Fig. 4f, Supplementary Fig. 4f). Among many targets, Epha2, a tyrosine receptor kinase, and Tbx3, a transcription factor, both activated by cMYC-driven signals in pre-pregnancy CAGMYC MECs, failed to accumulate H3K27ac signals and cMYC DNA occupancy in post-pregnancy CAGMYC MECs (Fig.4f, g and Supplementary Fig. 4f). Given that deletion of Epha2 in mouse models of mammary oncogenesis impacted tumor development and progression, this could represent a mechanism by which post-pregnancy CAGMYC MECs resist to *cMYC* overexpression [49].

## Discussion

Collectively, these results reveal a complex gene regulatory network supported by *cMYC* overexpression during mammary ductal hyperplasia, and demonstrate that such programs are not activated in post-pregnancy CAGMYC MECs. Our epigenetic and transcriptomic analyses corroborate our tissue morphology and cellular-based analyses, showing that *cMYC* overexpression failed to drive oncogenesis in post-pregnancy MECs.

Previous studies have characterized the altered molecular states of rodent and human post-pregnancy MECs, revealing an array of factors and signaling networks featured in post-pregnancy tissue, and providing insights into their potential contribution to breast cancer risk [50–53]. In our study, we have used a similar approach, but instead focused on the active regulatory epigenetic landscape of pre- and post-pregnancy murine MECs, with the goal of understanding its role in transcription activation, and in response to signals from consecutive pregnancies and initial stages of mammary oncogenesis.

Interestingly, the expansion of the pregnancy-induced active enhancer landscape minimally recapitulated the transcriptional output of post-pregnancy MECs during tissue homeostasis, as demonstrated by our gene expression analyses (Fig.1). This observation could suggest that the chromatin state captured in our analysis, defined by the activation mark H3K27ac alone, does not discriminate enhancer regions that were once highly active (during pregnancy) and are now in a less active/resting state, poised to respond to future pregnancy signals [54]. It is also possible that the abundance of specific TFs and epigenetic factors fluctuate across non-pregnancy, pregnancy, and post-pregnancy states, guiding chromatin remodeling and gene expression control. As there is a lack of information about how post-pregnancy cells reorganize their epigenome and the factors driving such changes, we have focused our analyses on the dynamics of enhancer activation and gene regulation in response to pregnancy and the early stages of oncogenesis, illustrating the complexity of the networks involved in these events.

Our epigenomic and transcriptomic analyses utilized a cell isolation strategy previously applied to isolate defined MEC populations [55, 56]. However, we cannot exclude that, after pregnancy, these cell-surface markers recognize a more diverse cell population compared to those existing prior to pregnancy. In fact, previous studies have reported the appearance of parity-induced mammary epithelial cells (Pi-MECs) [5, 57–61], suggesting an increased cellular heterogeneity after pregnancy. In agreement, recent single-cell RNA-seq analyses demonstrated MEC population alterations throughout gestation, lactation and involution stages of mammary gland development [62]. Our previous DNA methylation analyses of several mammary cell types from fully involuted, post-pregnancy mammary glands, demonstrated that alterations to the epigenome were, to some extent, shared by most mammary cell types [12], thus suggesting that, even in the case of population variation and increased heterogeneity, pregnancy-induced epigenomic alterations reach across mammary epithelium cell populations.

Our present study also revealed that post-pregnancy MECs are able to resist the oncogenic signals of *cMYC* overexpression, even in transplantation assays into pregnancy naïve environment. But how can pregnancy decrease mammary oncogenesis? Previous studies established that both a full pregnancy cycle (gestation, lactation, and involution), and an induced pseudo-pregnancy decreased the frequency of mammary tumors in several transgenic mouse strains, including those lacking p53 expression [20], those from chemically-induced mammary tumorigenesis [14, 21], and those accompanied by MMTV-driven *cMYC* overexpression [22]. Using an inducible system for controllable, *cMYC*-driven initiation of mammary oncogenesis, we lend further support the notion that post-pregnancy MECs resist the early stages of oncogenesis. We defined the effects of *cMYC* overexpression at both the level of tissues/cells, and at the level of the epigenome and transcriptome of MECs, showing that the cMYC cancer-driven programs were blocked in post-pregnancy CAGMYC MECs.

Previous studies have reported that pro-oncogenic signals which are naturally present immediately after pregnancy and during mammary-gland involution, increase tissue remodeling and activation of inflammatory signals that may support tumor development [63–65]. Thus, it is possible that during involution, *cMYC* overexpression, along with tissue remodeling and immune infiltration, has a different effect on the mammary epigenome and on mammary oncogenesis than those observed in fully involuted MECs (Fig.3, 4). Future studies mapping cMYC driven epigenomic alterations in CAGMYC MECs during involution will be important to distinguish the pregnancy-dependent signals driving either increased or decreased mammary tumorigenesis.

It is important to note that signals driving early oncogenesis may also differ substantially from those present after disease establishment. Our analysis of pre-pregnancy CAGMYC MECs suggests that transcriptional programs and the epigenome are differentially regulated across the *cMYC* overexpression timeline, supporting this notion. In fact, it has been previously shown that longer exposure to *cMYC* overexpression resulted in substantial reprogramming of breast epithelial cellular identity [66]. Our analysis did not address the effects of sustained *cMYC* overexpression on mammary tumor development, and instead focused on those that followed the progression from murine MEC normal cellular state into early onset of mammary oncogenesis, a largely unexplored field which warrants relevant hypotheses to understand epigenetic alterations and oncogenesis progression.

Our transplantation and organoid experiments were designed to define cell-autonomous features of post-pregnancy MECs in response to *cMYC* overexpression. However, it is possible that pregnancy induces alterations to the mammary gland stroma and/or its immune composition. Gene ontology analysis of H3K27ac signals specific to post-pregnancy CAGMYC MECs (Supplementary Fig. 4e) suggest activation of signals that may influence the extra cellular matrix (ECM). These signals likely arise from the post-pregnancy MECs themselves, further supporting the hypothesis that pregnancy-induced cell-autonomous signals that alter the early stages of oncogenesis, are perhaps accompanied by stromal alterations. It has also been suggested that pregnancy-induced alterations to the mammary gland ECM play a role in preventing the progression of established cancer cells [67]. Furthermore, alterations to immune composition after pregnancy have also been suggested to influence mammary tumor progression [68]. Thus, analyses exploring the correlation among stroma, immune system and the MEC epigenome need to be carried out to define their specific roles in blocking the early onset of oncogenesis in a post-pregnancy-dependent fashion.

Lastly, our data revealed the stability of the pregnancy-induced epigenome both in *in vivo* and *in vitro* pregnancy-naïve environments, observations that point to the cell-autonomous nature of the altered epigenome. Such *in vivo* and *in vitro* strategies will be required to monitor the detailed dynamics of enhancer activity and transcription regulation in a pregnancy-hormone fashion. In addition, 3D organoid cultures represent an ideal experimental system that should allow one to mechanistically dissect the signals that maintain and activate the epigenetic memory of pregnancy. Ultimately, such strategies may allow to understand enhancer activation and transcription regulation in post-pregnancy tissues obtained from women with various reproductive histories, and to study the effects of pregnancy-induced developmental dynamics on cancer predisposition.

## Methods

### Mouse lines

Balb/C and C57/BL6 female mice were purchased from Charles River. CAGs-rtTA3 mice (B6N.FVB(Cg)-Tg(CAG-rtTA3)4288Slowe/J, the Jackson Laboratory) and tetO-MYC mice (FVB/N-Tg(tetO-MYC)36aBop/J, the Jackson Laboratory) crossed for the establishment of CAGMYC transgenic mouse strain (129/C57BL6 background). All experiments were performed in agreement with approved CSHL Institutional Animal Care and Use Committee (IACUC).

### Mammary gland isolation

Mammary glands were harvested and processed as previously reported [56] and fully described in Supplemental Experimental Procedures. In short, mammary glands were harvested and into a single cell suspension. Epithelial cells were separated from immune cells (CD45+), red blood cells (TER119+) and endothelial cells (CD31+) using depletion antibodies and MACS magnetic column (Miltenyi Biotech). Lineage depleted (LIN-) epithelial cells were utilized for FACS-sorting, fat-pad transplantation assay, qPCR analysis and Western blot as described in Supplemental Experimental Procedures.

### Illumina library preparation and NextGen sequencing

FACS-isolated MECs were utilized for the preparation of NextGen Illumina libraries, as described in Supplemental Experimental Procedures.

### Histological analysis

Tissue histology and Immunofluorescence staining (IF) were performed as described in Supplemental Experimental Procedures.

## Author contributions

C.O.D.S. designed and supervised the research;; C.O.D.S, M.A.M., and M.J.F. wrote the manuscript. M.A.M. performed bioinformatics analysis;; M.J.F., C.C., S.L.C., S.T.Y., M.C., and W.D.F. performed experiments and analyzed results;; J.E.W. supported histology analysis.

## Acknowledgements

This work was performed with assistance from the CSHL Flow Cytometry Shared Resources, CSHL Animal Facility, the CSHL NextGen Sequencing Shared Resources, and the CSHL Histology Shared Resource, which are supported by the Cancer Center Support Grant 5P30CA045508. This work was supported by the Rita Allen Scholar Award, the V-foundation Scholar Award, the AACR-Breast Cancer Research Foundation Award and the Pershing Square Sohn Prize for Cancer Research (C.O.D.S.).

## Competing interests

The authors have no competing interests to disclose.

## Supplementary Information

### Materials and Methods

#### Antibodies

All antibodies were purchased from eBioscience, BioLegend or Abcam and used without further purification. Antibodies for lineage depletion: biotinylated anti-CD45 (eBioscience, #13-0451-85), biotinylated anti-CD31 (eBioscience, #13-0311-85), and biotinylated anti-Ter119 (eBioscience, #13-5921-85). Antibodies for flow cytometry: eFluor 450 conjugated anti-CD24 (eBioscience, #48-0242-82), PE-Cy7 conjugated anti-CD29 (eBioscience, #25-0291-82), PE conjugated anti-CD61 (BioLegend, #104307), APC conjugated anti-CD133 (BioLegend, #141208), PerCP-Cy5.5 conjugated anti-CD1d (BioLegend, #123514), 7-AAD viability staining solution (BioLegend, #420404). Antibodies for negative controls: FITC conjugated rat IgG (eBioscience, #11-4811-85), PE conjugated rabbit IgG (eBioscience, #12-4815-82), APC conjugated mouse IgG (eBioscience, #17-4210-82), PerCP-Cy5.5 conjugated mouse IgG (BioLegend, #405314), eFluor 450 conjugated mouse IgG (eBioscience, #48-4015-82) and PE-Cy7 conjugated mouse IgG (BioLegend, #405315). Antibody for MaSC enrichment: biotinylated anti-CD1d (BioLegend, #123505). Antibodies for immunostaining (IF): Alexa Fluor 488 conjugated anti-keratin 5 (Abcam, #ab193894), Alexa Fluor 405 conjugated anti-keratin 8/18 (Abcam, #ab210139), Alexa Fluor 647 conjugated anti-cMYC (Abcam, #ab190560). Antibodies for Western Blot: anti-cMYC antibody (Y69, Abcam, #ab19372), anti-GAPDH (SCBT, #sc-365062), goat anti-rabbit IgG HRP (Abcam, #ab6721) and goat anti-mouse IgG HRP (Abcam, #ab97051).

#### Flow cytometry

Mammary epithelial cells (MECs) were sorted using a FACS ARIAII SORP (BD Bioscience). For cell analysis, Dual Fortessa II cell analyzer (BD Bioscience) or MACSQuant (Miltenyi Biotech) were utilized. Data analysis was performed using FloJo (Tree Star).

#### Doxycycline (DOX) treatment

Doxycycline was purchased from Takara Bio USA, Inc. (#631311) and sucrose was purchased from Sigma (S7903). DOX drinking solution (1 mg/mL) was prepared using sterile 1% sucrose water. CAGMYC female mice and all control mice were given DOX water for 5 days or otherwise specified.

#### Post pseudo-pregnancy analysis

Balb/C (Fig.1 and Fig.3), CAG-only transgenic (Fig.3) and C57/BL6 (Fig.3) female mice were implanted with slow-release estrogen and progesterone pellets (17*ββ*-Estradiol (0.5 mg/pellet) + Progesterone (10 mg/pellet) – Innovative Research of America), or placebo control pellets as previously described [1]. Mammary glands were harvested ∼60 days post pellet implantation (∼20 days pseudo-pregnancy + 40 days forced involution) for downstream analysis.

#### Mammary gland isolation

In short, mammary glands were harvested, minced and incubated for ∼2 hours with 1x Collagenase/Hyaluronidase (10x solution, Stem Cell Technology) in RPMI 1640 GlutaMAX supplemented with 5% FBS. Digested mammary gland fragments were washed with cold HBSS supplemented with 5% FBS, followed by incubation with TrypLE Express (Thermo Fisher, #12604-013) and an additional HBSS wash. Cells were incubated with 2 mL of Dispase (Stem Cell Technology) supplemented with 40 µL DNAse I (Sigma, #D4263) for 2 minutes and then filtered through a 100μm Cell Strainer (BD Falcon, #352360). The single cell suspension was incubated with lineage depletion antibodies and loaded onto MACS magnetic column (Miltenyi Biotech). Flow-through cells (epithelial cells) were utilized in tissue culture, protein isolation, or flow cytometry analysis.

#### Histological analysis

For histological analysis, the left inguinal mammary gland was harvested and fixed in 4% PFA overnight prior to paraffin-embedding. For conventional histological analysis, mammary gland tissue slides were stained with Hematoxylin and Eosin (H&E). Images were acquired using Aperio ePathology (Leica Biosystems) slide scanner in 40X lenses. For Immunofluorescence staining (IF), paraffin-embedded mammary gland sections were deparaffinized in Xylene (Sigma, #534056) and rehydrated, followed by antigen retrieval in Trilogy (Cell Marque, 920P-10). Tissue was washed in 1x PBS (phosphate-buffered saline) for 1 min then blocked with blocking solution (10 mM Tris-HCl pH 7.4, 100 mM MgCl_2_, 0.5% Tween 20, 10% FBS, 5% goat serum) for 4 hours in a humidified chamber. Sections were stained with the appropriate conjugated primary antibodies in blocking solution for 16 hours at 4°C. After subsequent washings with 1x PBS and blocking solution, slides were mounted in ProLong with DAPI (Invitrogen). Cell visualization and image collection was performed on a Zeiss LSM780 confocal laser-scanning microscope utilizing Zen lite software (Zeiss). Magnification: 20X. Statistically significant differences for ductal quantification were considered with Student t-test *p*-value lower than 0.05 (p<0.05).

#### Mammary fat pad transplantation

MaSCs from pre- and post-pregnancy Balb/C female mice (Fig.1) or pre- and post-pregnancy CAGMYC female mice (Fig.3) were harvested and digested as previously described [2]. Lineage depleted MECs were incubated with biotinylated anti-CD1d to allow for MaSCs enrichment, followed by MACS magnetic column isolation. CD1d-enriched MaSCs were ressuspended with 50% growth factor reduced matrigel solution (BD Biosciences) and injected into the cleared fat-pad of the inguinal mammary gland of Balb/C female mice (Fig.1) or CAG-only transgenic female mice (Fig.3). Transplanted mammary glands were then harvested for downstream analysis and ductal quantification. Statistically significant differences were considered with Student t-test *p-*value lower than 0.05 (p<0.05).

#### RT-PCR and qPCR analysis

FACS-sorted MECs were homogenized in TRIzol (Thermo Fisher Scientific, #15596018) for RNA extraction. Double stranded cDNA was synthesized from purified RNA using SuperScript III Reverse Transcriptase (Thermo Scientific). For qPCR analysis, qPCR primers (Sup. Table 1) were utilized for quantitative PCR reactions on QuantStudio 6 real time PCR system (Thermo Fisher) and quantification results were analyzed using the delta delta CT method. For ChIP-qPCR, FACS-sorted MECs were used for chromatin pulldowns using antibodies specific for cMYC (Abcam, #ab32072) and rabbit IgG (Millipore, #12-371). ChIP samples were analyzed by quantitative PCR reactions on a Quantstudio 6 real time PCR system (Thermo Fisher), utilizing primers designed across the transcription start site of the genes Epha2 and Tbx3 (Sup. Table 2). Results are displayed as fraction of Input (non-immunoprecipitated DNA). n=2. Error bars represent standard variation of mean values. Statistically significant differences were considered with Student t-test *p-*value lower than 0.05 (p<0.05).

#### Mammary Organoid Culture

Mammary glands were dissected from nulliparous and parous Balb/C female mice (Fig.1) or CAGMYC female mice (Fig.3), minced and digested for ∼40 minutes in Collagenase A, type IV solution, following a series of centrifugations to enrich for mammary organoids [3]. Organoids were cultured with either Essential media (Advanced DMEM/F12, supplemented with ITS (Insulin/Transferrin/Sodium selenite, Gibco, #41400-045, and FGF-2 (PeproTech, #450-33) or Complete media (Advanced DMEM/F12, supplemented with ITS (Insulin/Transferrin/Sodium selenite, Gibco, #41400-045, FGF-2 (PeproTech, #450-33), Prolactin (Sigma, #L4021), 17βEstradiol (Sigma #E2758) and Progesterone (Sigma, #P8783). Medium was changed every 3 days and organoids were passaged every 1-2 weeks. For experiments described on Figure 3, pre- and post-pregnancy CAGMYC organoids were treated with doxycycline (DOX, 0.1 mg/mL). Cell visualization and image collection was performed on a Nikon Eclipse TI microscope utilizing NIS-Elements BR software (Nikon).

#### Western blot

MECs from untreated or DOX-treated (DD5) pre- or post-pregnancy CAGMYC transgenic mice, were isolated and homogenized in 1x Laemmli sample buffer (Bio-Rad). Samples were loaded into home-made 10% SDS-Page gel and transferred overnight to PVDF membrane using wet-transfer apparatus (Bio-Rad). Membranes were blocked with 1% BSA solution and incubated overnight with a diluted solution of primary antibody, followed by incubation with HRP-conjugated antibody for 40 minutes. HRP signal was developed with Luminata Crescendo Western HRP substrate (Millipore) in autoradiography film (Lab Scientific, #XARALF2025). Developed films were scanned on Epson Perfection 2450 photo scanner.

#### MEC preparation for NextGen sequencing

Mammary glands from several Balb/C female mice (Fig.1, ∼20 weeks old) or CAGMYC transgenic female mice (Fig.2, ∼20 weeks old) were pulled, according to the following experimental design: Balb/C female mice pre-pregnancy (5 mice), post-pregnancy (5 mice), first pregnancy cycle at day 6 (D6) and day 12 (D12) (3 mice per time point), second pregnancy cycle at day 6 (D6) and day 12 (D12) (3 mice per time point), and DOX–treated CAGMYC pre-pregnancy at day 2 (DD2) and at day 5 (DD5) (2 mice per treatment). Mammary glands from individual CAGMYC transgenic female mice (Fig.4, total of 2 females per experimental group, ∼20 weeks old) were prepared according to the following experimental design: DOX–treated CAGMYC pre-pregnancy at day 5 and DOX–treated CAGMYC post-pregnancy at day 5 (DD5). Mammary glands were enzymatically digested, lineage depleted, and stained with antibody cocktail for the isolation of luminal and myoepithelial MECs (CD24, CD29), with the exception of data presented on Fig.2, in which total MECs were utilized.

#### RNA-seq library preparation and analysis

Total MECs or FACS-sorted MECs were collected and homogenized in TRIzol (Thermo Fisher Scientific, #15596018) for RNA extraction. Double stranded cDNA synthesis and Illumina libraries were prepared utilizing the Ovation RNA-seq system (V2) (Nugen Technologies, #7102-32). RNA-seq libraries were prepared utilizing the Ovation ultralow DR multiplex system (Nugen Technologies, #0331-32). Each library (n=2 per experimental condition) was barcoded with Illumina True-seq adaptors to allow sample multiplexing, followed by sequencing on an Illumina NextSeq500, 76 bp single-end run. We used STAR [4] for mapping reads. We used DESeq [5] to assess changes in expression levels simultaneously across multiple conditions and in multi-factor experimental designs, incorporating information from multiple replicates (2 independent experiments per cell type). Gene Set Enrichment Analysis (GSEA) was used for global analysis of differentially expressed genes [6, 7]. Batch effect normalization of CAGMYC RNA-seq and publicly available microarray data was performed using the ComBat function from the ‘sva’ R package [8]. Further downstream analyses were performed using various R packages, including gplots for heatmaps [9], base R [10], and ggplots2 for other visualizations [11].

#### ChIP-seq library preparation and analysis

FACS-sorted MECs were used for chromatin pulldowns using antibody specific for H3K27ac histone marks (Abcam, #ab4729). ChIP samples were amplified and barcoded using Clontech DNA Smart ChIP-Seq kit (Clontech, #634866) in accordance with the manufacturer’s instructions, then pooled for sequencing on an Illumina NextSeq500, 76 bp single-end run (n=2 per experimental condition). Reads were mapped to the indexed mm9 genome using bowtie2 short-read aligner tool [12], using default settings. MACS2 peak-calling program [13] was used to identify enriched genomic regions in this data by comparing the pulldown ChIP data to the control (Input) data using a tag size of 25 bp and a q-value cutoff of 1.00^−­2^. The ROSE (Rank Order of Super-Enhancers) algorithm [14] was used to identify enhancers and super-enhancers throughout the genome using ChIP-seq BAM files and GFF files. GFF files were made from MACS-called BED files. ROSE analysis was performed using default conditions, which included a 2.5 Kb exclusion zone surrounding the TSS and a 12.5 Kb stitching distance. The UCSC genome browser was used to analyze enhancers for overlap via the table browser intersect function. Any overlap between regions (enhancers) was considered “shared”, whereas and no overlap between regions (enhancers) defined the regions as ONLY being in one sample. Identification of genes closest to these differentially called enhancer regions was preformed using Genomic Regions Enrichment of Annotations Tool (GREAT) [15]. Homer’s annotatePeaks.pl script was run with default settings in order to define genomic distributions. The heatmap of the peak intensities was produced using the plotHeatmap tool in deepTools2 [16]. Peak visualizations were generated using the UCSC Genome Browser [17].

#### ChIP-qPCR

FACS-sorted MECs were used for chromatin pulldowns using antibodies specific for cMYC (Abcam, #ab32072) and rabbit IgG (Millipore, #12-371). ChIP samples were analyzed by quantitative PCR reactions on a Quantstudio 6 real time PCR system (Thermo Fisher), utilizing primers designed across the transcription start site of the genes Epha2 and Tbx3 (Sup. Table 2). Results are displayed as fraction of Input (non-immunoprecipitated DNA). n=2.

**Supplementary Figure 1.**
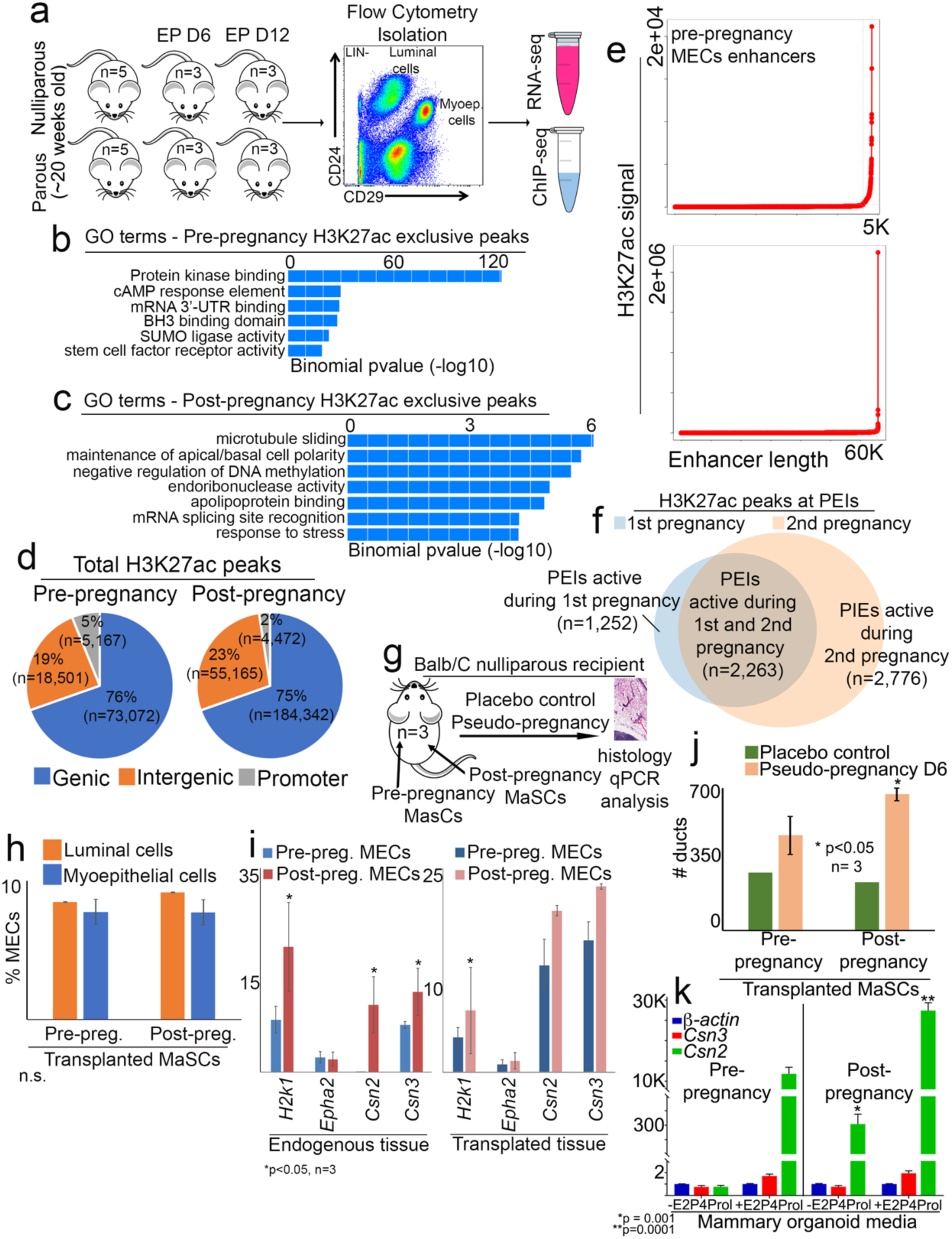
Characterization of epigenetic, transcriptomic and developmental state of pre- and post-pregnancy MECs. **(a)** Scheme of experimental strategy for epigenetic and transcription analysis of MECs spanning diverse development stages. (**b,c**) Gene Ontology term analysis of H3K27ac peaks exclusive to pre-pregnancy MECS (**b**), or exclusive to post-pregnancy MECS (**c**). (**d**) Classification and fraction of H3K27ac in pre- and post-pregnancy MECs according to their genomic location (genic, intergenic or promoter region). (**e**) Classification of H3K27ac peaks from pre- and post-pregnancy MECs into enhancer regions. (**f**) Distribution of distribution according H3K27ac intensity (defined as active regions) in MECs during first and second pregnancy. (**g**) Scheme of experimental design utilized for pre- and post-pregnancy MaSCs transplantation into nulliparous Balb/C female recipient, and analysis of tissue development in response to pseudo-pregnancy. (**h**) FACS quantification of viable (7AAD^−­^), lineage negative (Lin^−­^) luminal and myoepithelial MECs from mammary glands transplanted with pre- or post-pregnancy MaSCs (8 weeks post-transplantation). n=3. n.s.= not significant. (**i**) qPCR quantification of genes identified as up-regulated after pregnancy in MECs isolated from mammary glands transplanted with either pre- or post-pregnancy MaSCs. n= 3. *p<0.05. (**j**) Quantification of ductal structures from mammary glands transplanted with either pre- or post-pregnancy MaSCs at day 6 of pseudo-pregnancy or placebo control. Bars indicate mean number of ducts. Error bars indicate standard deviation amongst samples from same experimental group. n= 3. *p<0.05. (k) qPCR quantification of genes identified as up-regulated by MECs during second pregnancy in pre- and post-pregnancy mammary organoid cultures, grown without (-E2P4Prol) and with (+E2P4Prol) pregnancy hormones. n=3. *p=0.001 – differences between pre- and post-pregnancy mammary organoids grown without pregnancy hormones (-E2P4Prol);; *p=0.0001 - differences between pre- and post-pregnancy mammary organoids grown with pregnancy hormones (+E2P4Prol).

**Supplementary Figure 2.**
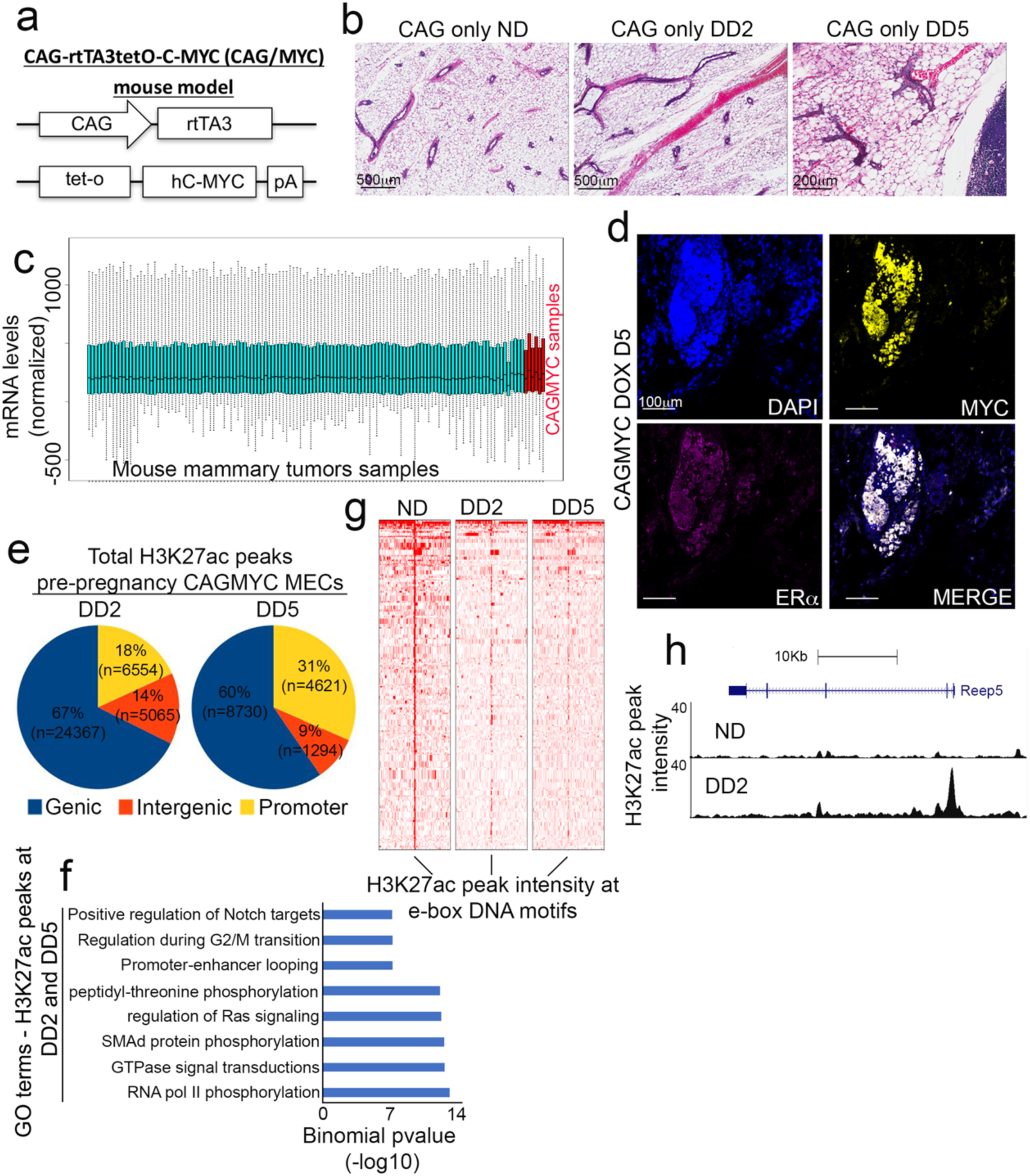
Inducible, short-term, *cMYC* overexpression drives early onset oncogenesis in mammary glands from nulliparous female mice. **(a)** Diagrammatic representation of the transgenic mouse strain CAGMYC showing alleles for overexpression of human *cMYC*, driven by the CAG promoter, under the inducible control of the Tet operator-repressor system. **(b)** H&E stained mammary gland images from nulliparous CAGMYC female mice showing the effects of sustained DOX-treatment on gland morphology in the absence of *cMYC* transgenic allele (CAG-only). Images acquired on Aperio ePathology (Leica Biosystems) slide scanner. **(c)** Normalization of publicly available gene expression datasets from transgenic mice of mammary tumor development and gene expression datasets generated from MECs harvested from DOX-treated CAGMYC mammary glands. **(d)** Immunofluorescence images of mammary glands from nulliparous CAGMYC female mice treated with DOX for 5 days (DD5), visualizing DAPI (blue), ER*αα* (magenta) and *cMYC* (yellow). Images captured on Zeiss LSM780 laser scanning microscope and processed on Zen lite (Zeiss). Scale: 100 µm. **(e)** Classification and percentage of H3K27ac peaks from pre-pregnancy CAGMYC MECs treated with DOX for 2 days (DD2, left) and 5 days (DD5, right), according to their genomic distribution. **(f)** Gene Ontology term analysis of H3K27ac peaks shared between pre-pregnancy CAGMYC MECs treated with DOX for 2 days (DD2) and 5 days (DD5). **(g)** Density plot showing the intensity of H3K27ac peaks at e-box DNA binding motifs in pre-pregnancy MECs from no DOX (ND) and DOX-treated MECs (DD2, DD5). **(h)** Genome Browser tracks for Reep5 gene showing differential H3K27ac distributions in CAGMYC MECs upon *cMYC* overexpression.

**Supplementary Figure 3.**
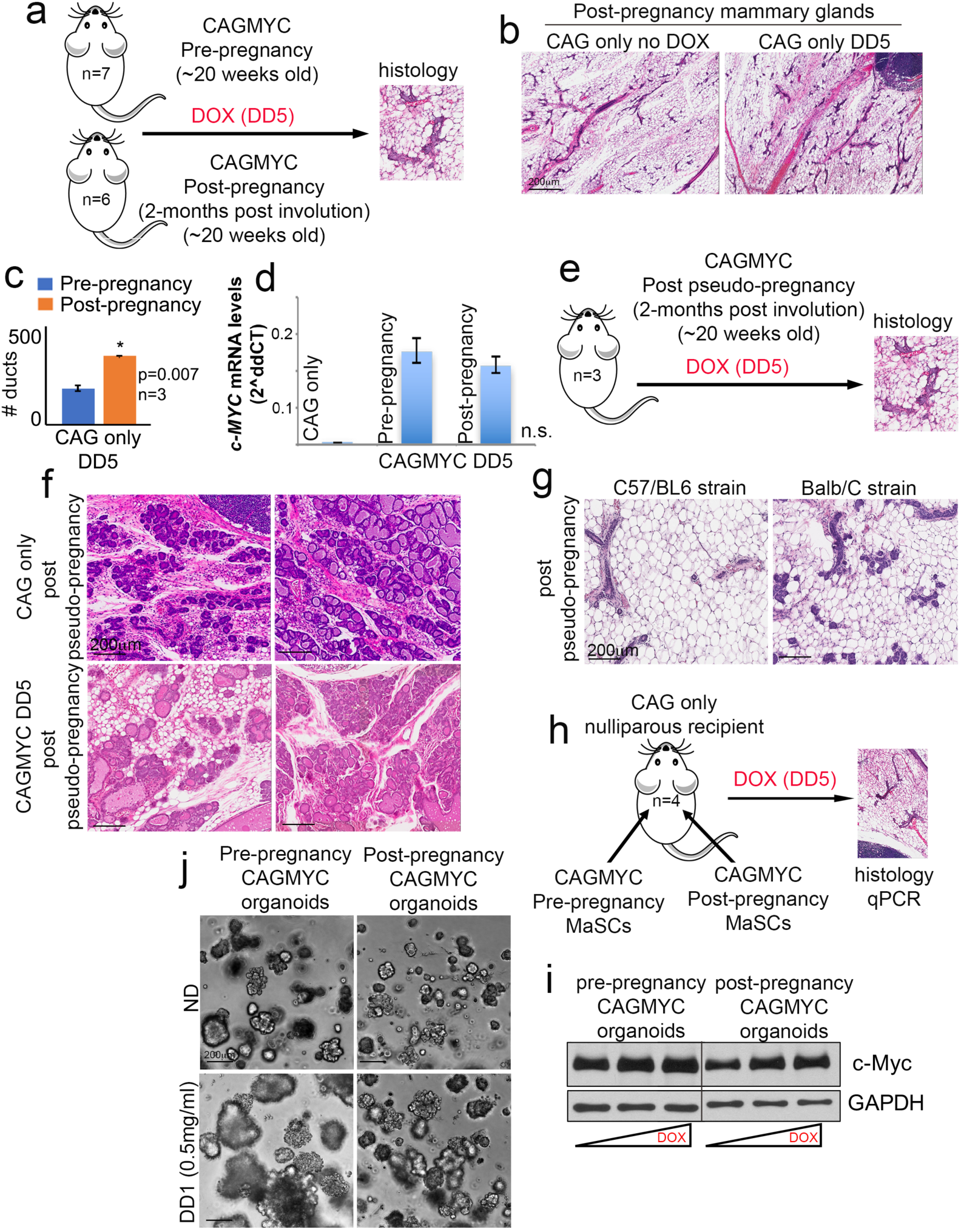
Defining the effects of short-term, *cMYC* overexpression on early onset oncogenesis in mammary glands from post-pregnancy female mice. **(a)** Scheme of experimental strategy for inducing *cMYC* overexpression in nulliparous and parous CAGMYC female mice by DOX treatment (DD5). **(b)** H&E stained mammary gland images from parous transgenic female mice showing the effects of DOX-treatment on gland morphology in the absence of *cMYC* transgenic allele (CAG-only). Images acquired on Aperio ePathology (Leica Biosystems) slide scanner. **(c)** Quantification of ductal structures in H&E-stained mammary glands from nulliparous and parous CAG only transgenic female mice after DOX treatment (DD5). n=3. Bars indicate mean number of ducts. Error bars indicate standard deviation amongst samples of same experimental group, n=3. *pvalue= 0.007. **(d)** Quantification of human *cMYC* mRNA levels by qPCR in control (CAG only) and in pre- and post-pregnancy CAGMYC MECs treated with DOX for 5 days. Bars indicate mean log2 fold change. Error bars indicate standard deviation amongst samples of same experimental group. n=?. n.s.= not significant. **(e)** Scheme of experimental strategy for post pseudo-pregnancy analysis in CAGMYC female mice after DOX treatment (DD5). **(f)** H&E stained mammary gland images from CAG only (top panels) and CAGMYC female mice (bottom panels) after completion of a pseudo-pregnancy. Scale: 200 µm. **(g)** H&E stained mammary gland images from C57/BL6 (left) and Balb/C female mice after completion of a pseudo-pregnancy. Scale: 200 µm. **(h)** Scheme of experimental strategy for analysis of nulliparous CAG only female mice transplanted with pre- and post-pregnancy CAGMYC MaSCs, followed by DOX treatment (DD5). **(i)** Western blot analysis of *cMYC* and GAPDH levels in pre- and post-pregnancy CAGMYC organoid cultures treated with several concentrations of DOX (0.01 mg/mL, 0.1 mg/mL or 0.5 mg/mL). **(j)** Images of pre- and post-pregnancy CAGMYC organoid cultures grown for 24 hours with high concentration of DOX (0.5 mg/mL). Images captured on Nikon Eclipse TI microscope and processed on NIS-Elements BR (Nikon). Scale: 200 µm.

**Supplementary Figure 4.**
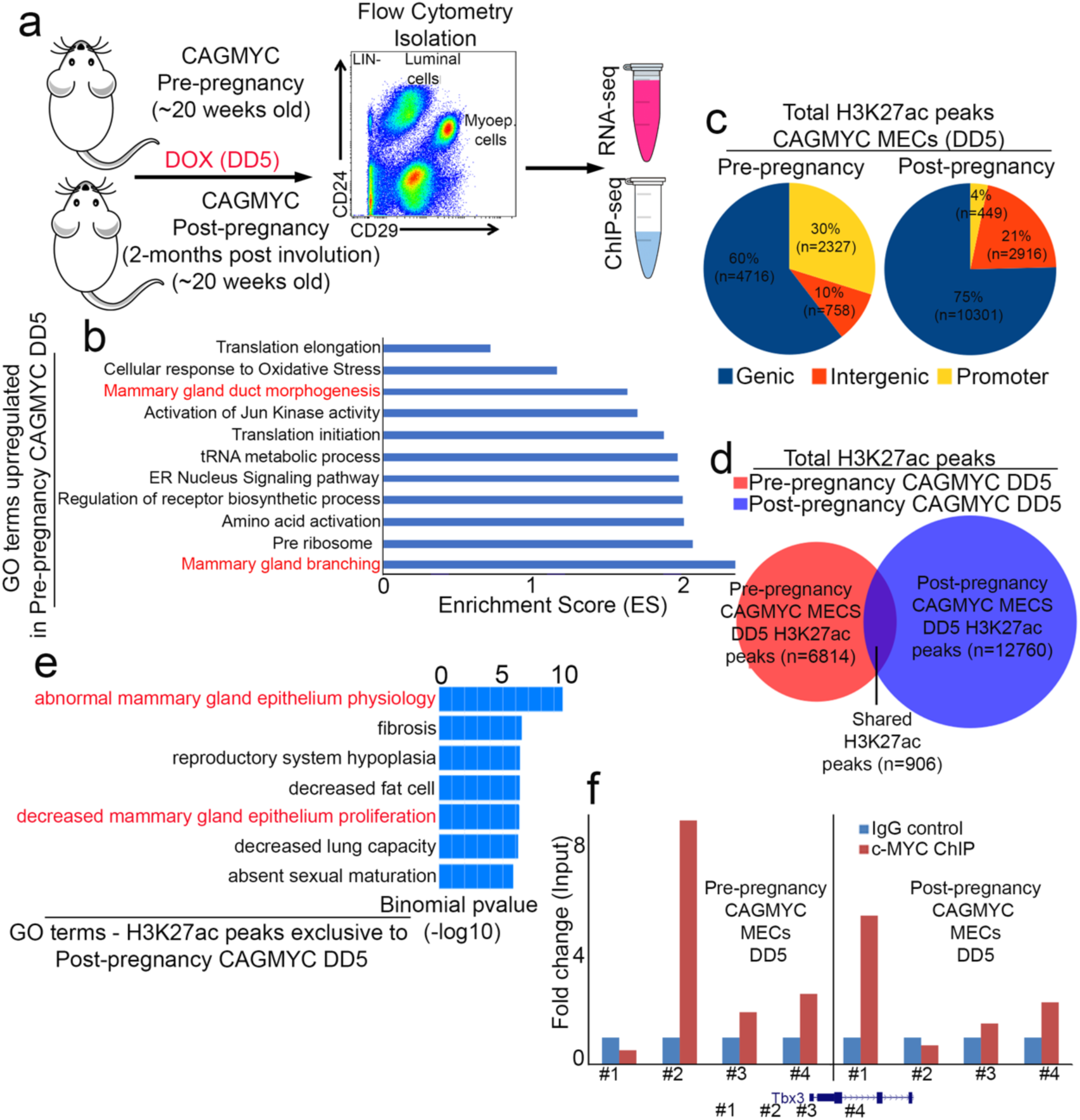
Epigenomic and gene expression changes induced by cMYC overexpression in post-pregnancy CAGMYC MECs. **(a)** Scheme of experimental strategy for epigenomic and transcriptomic analysis of isolated MECs from nulliparous and parous CAGMYC female mice after DOX treatment (DD5). (**b**) Gene ontology terms yielded from GSEA analysis for transcriptional programs differentially expressed in post-pregnancy CAGMYC MECs treated with DOX for 5 days. NES: Normalized Enrichment Score. pvalue <0.05. (**c**) Classification and percentage of H3K27ac peaks from pre- and post-pregnancy CAGMYC MECs treated with DOX for 5 days (DD5), according to their genomic distribution. (**d**) Venn diagram showing unique and shared H3K27ac peaks in pre- and post-pregnancy CAGMYC MECs treated with DOX for 5 days (DD5). (**e**) Gene Ontology term analysis of H3K27ac peaks exclusive to post-pregnancy CAGMYC MECs. (**f**) ChIP-qPCR quantification of cMYC occupancy at the *Tbx3* gene locus in pre- and post-pregnancy CAGMYC MECs treated with DOX for 5 days. Bars indicate mean fold change.

**Table S1.**
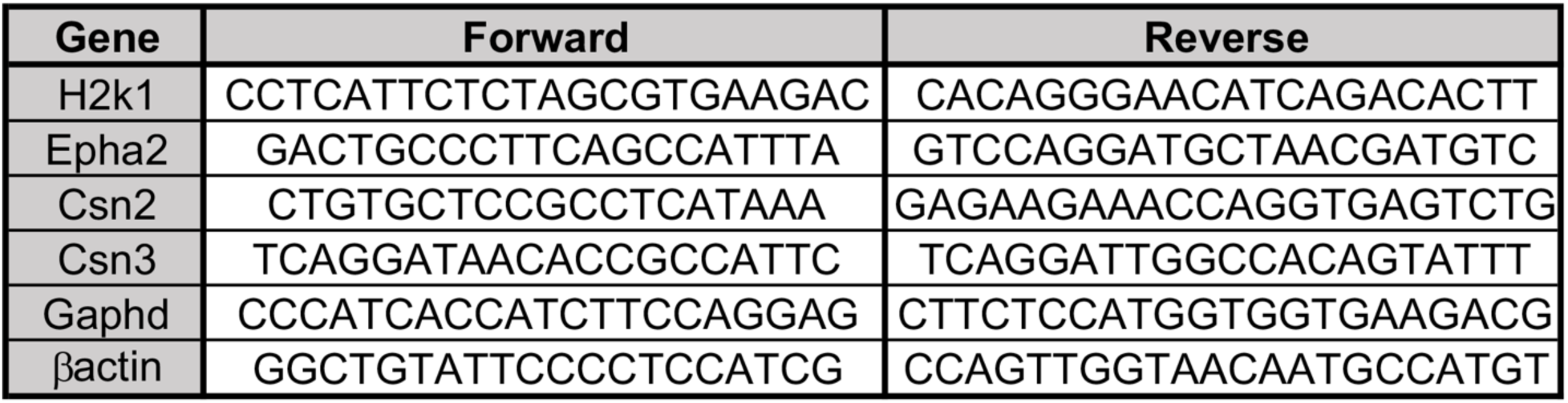
Primer list for genes utilized in qPCR analysis.

**Table S2.**
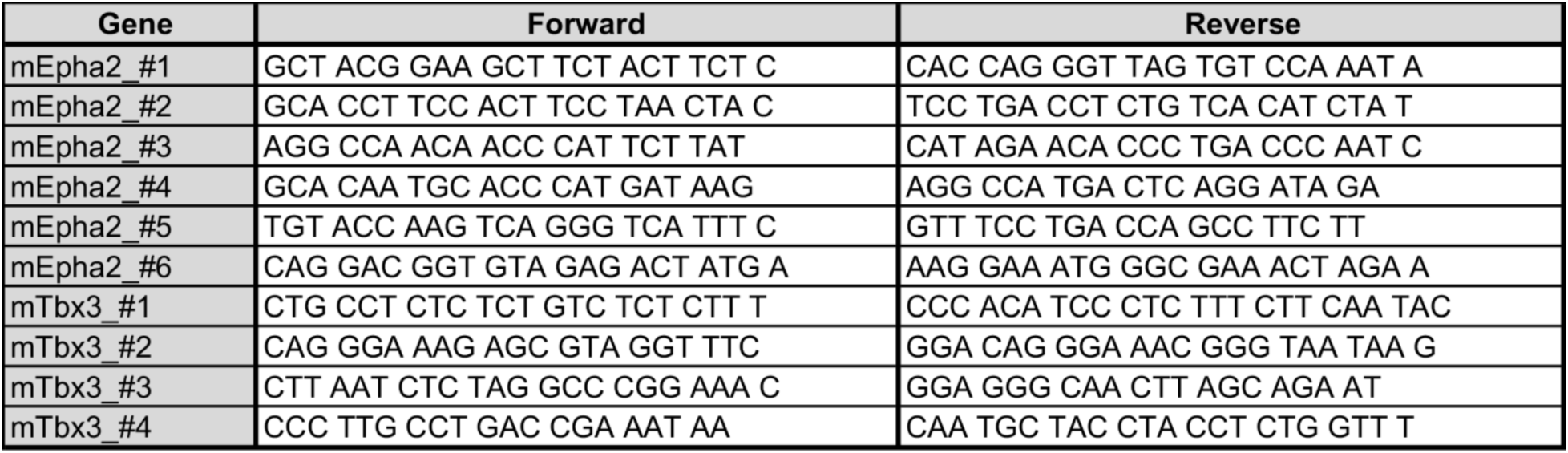
Primer list for genes utilized in ChIP-qPCR analysis.

